# Extreme suction attachment performance from specialised insects living in mountain streams (Diptera: Blephariceridae)

**DOI:** 10.1101/2020.09.30.320663

**Authors:** Victor Kang, Robin T. White, Simon Chen, Walter Federle

**Affiliations:** Department of Zoology, University of Cambridge, Cambridge, CB2 3EJ, United Kingdom; Carl Zeiss Research Microscopy Solutions, Pleasanton, California, 94588, United States

**Keywords:** Biomechanics, adhesion, underwater adhesion, aquatic invertebrates, surface roughness

## Abstract

Suction is widely used by animals for strong controllable underwater adhesion but is less well understood than adhesion of terrestrial climbing animals. Here we investigate the attachment of an aquatic insect larva (Blephariceridae), which clings to rocks in torrential streams using the only known muscle-actuated suction organs in insects. We measured their attachment forces on well-defined rough substrates and found their adhesion was much less reduced by micro-roughness than terrestrial climbing insects. *In vivo* visualisation of the suction organs in contact with microstructured substrates revealed that they can mould around large asperities to form a seal. Moreover, we showed that spine-like microtrichia on the organ are stiff cuticular structures that only make tip contact on smooth and microstructured substrates. Our results highlight the performance and versatility of blepharicerid suction organs and introduce a new study system to explore biological suction.

## Introduction

Of the approximately one million known species of insects, only 325 attach using muscle-controlled suction organs [1, 2]. These species belong to a single Dipteran family, the Blephariceridae, and their larvae and pupae develop on rocks in torrential alpine streams where flow-rates can exceed 3 ms^-1^ [3, 4]. Each blepharicerid larva has six ventral suction organs to attach to biofilm-covered rock surfaces, where it feeds on diatoms. Using its suction organs, the larva can locomote relatively quickly and possibly over long distances: blepharicerid larvae migrate from one stone to another to find the swiftest regions of the stream [5, 6]. Once development is complete, the winged adult emerges from its pupa, floats to the water surface, and immediately flies away to mate and lay eggs to begin the cycle anew [7, 8]

The remarkable morphology of blepharicerid suction organs is well-described [6, 9–11]. The organ superficially resembles a synthetic piston-pump, with a suction disc that interacts with the surface and creates a seal, a central piston and powerful piston muscles to manipulate the pressure, and a suction chamber with a thick cuticular wall to withstand low pressures during attachments. There are spine-like microstructures called microtrichia on the suction disc that contact glass surfaces and may increase resistance to shear forces. In addition, a dedicated detachment system allows the larva to rapidly detach its suction organ during locomotion [10].

While much is known about their morphology, the mechanisms involved in blepharicerid suction attachment are less well understood. Two studies to date have measured the attachment performance of blepharicerid larvae [12, 13], yet neither of them offers mechanistic insight into how their suction organs cope with different surface conditions to generate strong underwater attachments. The limited state of knowledge extends to biological suction in general: there are only a few well-studied animals (namely, remora fish, clingfish, octopus, and leeches) for which the function of specific structures in biological suction attachments has been experimentally demonstrated [14– 23]. This may be the consequence of mechanistic studies on biological adhesion focussing primarily on terrestrial climbing animals such as geckos, tree-frogs, insects, and spiders [24, 25], but it is still surprising as suction is one of the main strategies for strong and controllable underwater adhesion. Nevertheless, the fundamental principles derived from terrestrial animals have greatly expanded our knowledge on how to achieve and control adhesion in air. Likewise, additional mechanistic studies on biological suction can help to identify new strategies for generating and controlling underwater adhesion on different surface conditions.

Here we investigate the mechanisms underlying suction attachments of blepharicerid larvae. We first conduct a detailed morphological study of *Hapalothrix lugubris* (Blephariceridae) to provide new insights into structures that are relevant for suction attachments. To understand how well blepharicerid larvae attach to different surfaces, we quantify their performance on smooth, micro-rough, and coarse-rough surfaces. We examine the function of spine-like microtrichia through *in vivo* visualisation of the contact zone during attachments on smooth and microstructured substrates. Furthermore, we compare blepharicerid suction performance with that of a model terrestrial insect to investigate how two fundamentally distinct adhesive systems cope with surface roughness.

## Results

### Morphology of the suction attachment organ of *Hapalothrix lugubris*

*H*.*lugubris* larvae have six ventromedian suction organs with each organ comprising a suction disc, a central opening and a piston, a suction chamber surrounded by a thick-walled cuticular cuff, and a V-notch (Figure 1 and Video 1). The suction disc contacts the surface for attachment, and the piston and underlying piston muscles (Figure 1c & d) actively lower the pressure inside the suction chamber. Two apodemes attaching to the V-notch in *H. lugubris* mediate its muscle-controlled opening for rapid detachment of the suction organ (Figure 1e; see also [10]).

**Figure 1.**
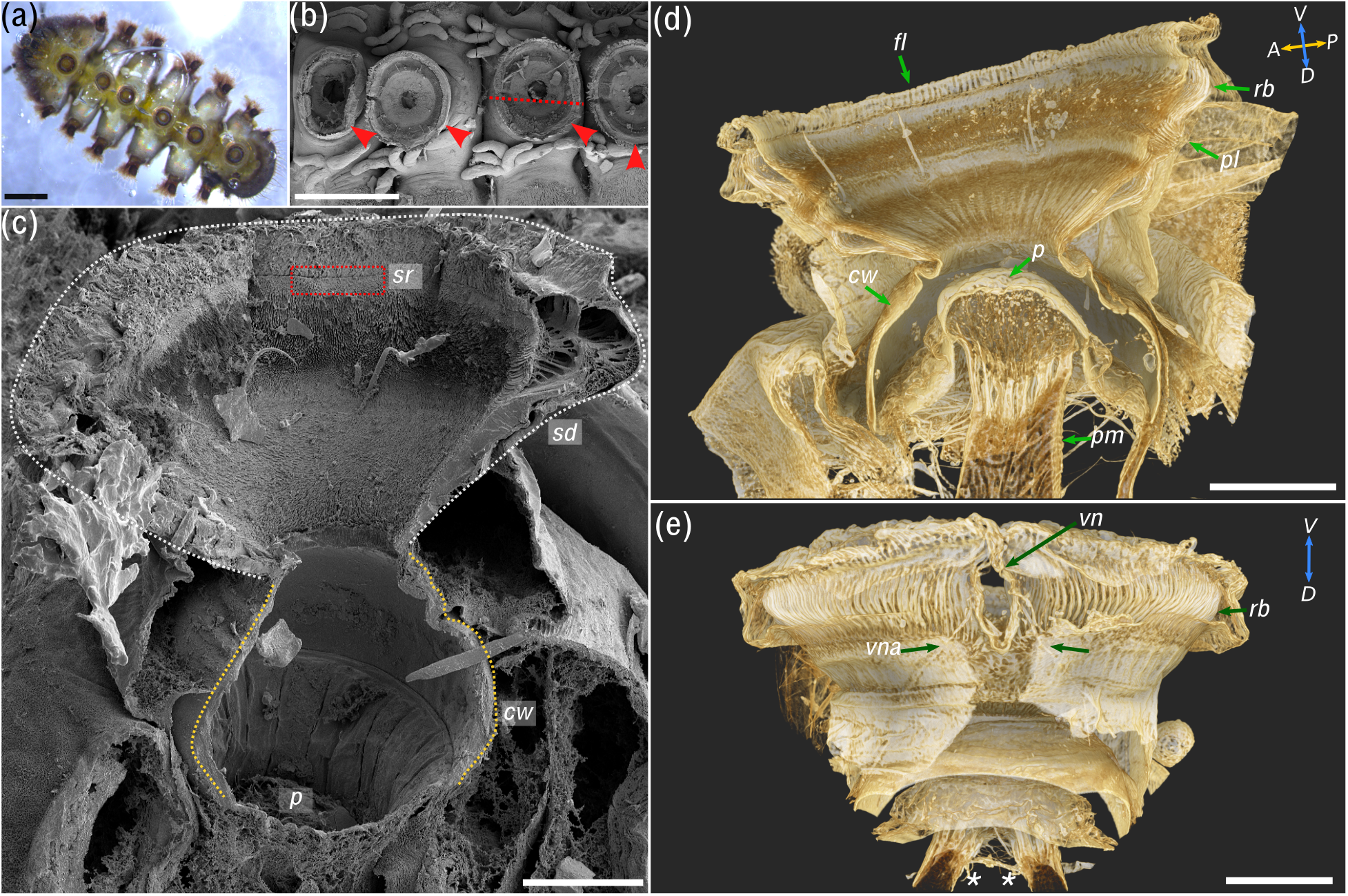
Overview of *Hapalothrix lugubris* the suction attachment organs. (a) Six ventromedian suction organs of a third instar *H. lugubris*. (b) Scanning electron micrograph of a flash-frozen and freeze-dried larva. Red arrows point to suction organs. Tracheal gills are visible on both sides of the organ. Dotted line shows approximate location of fracture line for (c). (c) Freeze-fracture reveals components important for attachment: suction disc (outlined in white dotted line, *sd*), sealing rim (red box, *sr*), cuff wall (yellow dotted line, *cw*), and the piston (*p*). The ventral surface of the suction disc is covered by a dense array of microtrichia (see Figure 2 for details). (d) **Micro-CT rendering of the organ highlights its internal organisation in 3D**. (d) Sagittal view, showing the following structures: outer radial beams (*rb*), palisade cell layer (*pl*), piston cone (*p*), and piston muscles (*pm*). The cuff wall (*cw*) encircles the suction cavity, and the outer fringe layer (*fl*) encircles the disc. *A*: anterior, *P*: posterior, *D*: dorsal, V: ventral. (e) View from anterior side, showing the V-notch (*vn*) and its pair of apodemes (*vna*) extending dorsally into the body. Outer cuticle has been digitally dissected to reveal the radial beams. Note the pair of piston muscles extending dorsally (*). Scale bars: (a) & (b) 500 µm; (c) 50 µm; (d) & (e) 100 µm.

The ventral disc surface of *H. lugubris* is covered in a dense array of microtrichia (Figures 1c & 3). The suction disc sealing rim, which seals the disc for suction attachment, closely resembles that of *L. cinerascens* [10] and comprises a dense array of upright rim microtrichia (Figure 1c). This is different to *L. cordata*, which has a distinct rim made up of a single row of flat rim microtrichia [10]. Going from the rim to the centre of the disc, the rim microtrichia transition into longer spine-like microtrichia (6.7 ± 0.5 µm in length; mean of means ± standard error of the mean; n=2 individuals), and then again to shorter microtrichia in the centre.

Multiple imaging techniques were used to gain additional insights into the ultrastructure and internal organisation of the suction disc: freeze-fracture scanning electron microscopy (SEM), 3D models using computed microtomography (micro-CT) data, and *in vivo* transmitted light microscopy (Figure 2). While internal fan-fibre networks underneath the outer regions of the suction disc have been mentioned previously [9] (Figure 2a), we discovered that each internal fibre leads to a single microtrichium (Figure 2b). Moreover, all the microtrichia that were fractured during sample preparation appeared to be solid (in-filled) cuticular structures (Figure 2c). The small internal fibres leading into the microtrichia branch out from thicker trunks originating from the ventral side of the outer radial beams (Figure 2b & d). The radial beams are also solid cuticular structures and alternate between a wide and a narrow beam (Figure 2d). The beams originate from the palisades, a radial zone consisting of dorsoventral cuticular rods (Figure 2e & f). There are 72 radial beams in a 90° segment of the disc, corresponding to 288 beams per disc (assuming no interruption from the V-notch) and a centre-to-centre spacing of around 4 µm or 1.3°.

**Figure 2.**
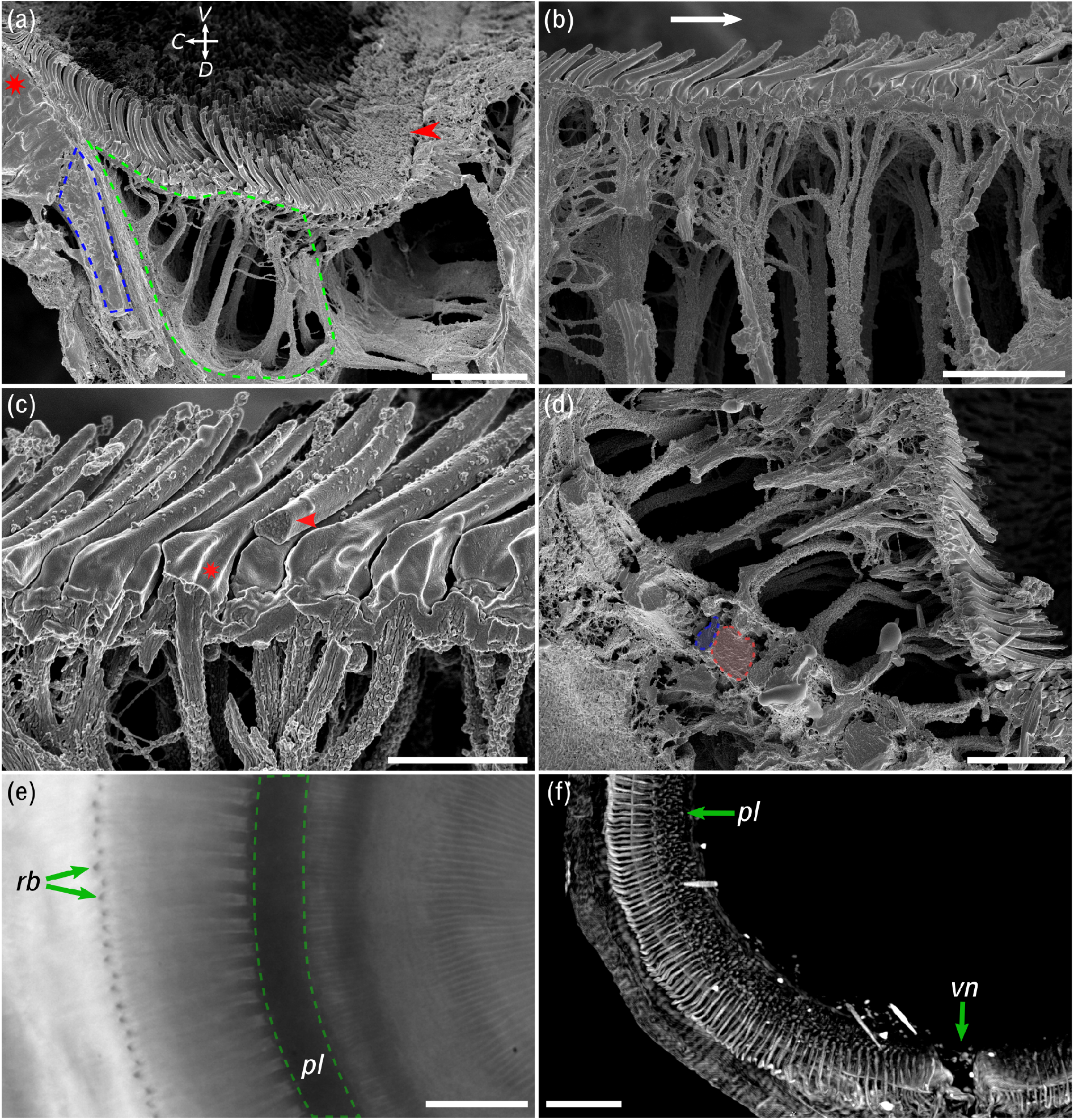
Ultrastructure of the suction disc. (a-d) Scanning electron micrographs of freeze-fractured suction organ cuticle (radial fracture plane). (a) Outer region of suction disc. The sealing rim and its short rim microtrichia are marked by a red arrowhead. Internal radial beams (highlighted in blue) originate from the palisade layer (red *). The fan-fibre space is highlighted in green. Note the spine-like microtrichia point towards the disc centre. *V*: ventral, *D*: dorsal, *C*: disc centre. (b-d) **Details of microtrichia in region of fan-fibre space**. (b) Each microtrichium connects to an internal fibre; these fine fibres represent the endings of thicker branched fibres originating from the radial beams. Arrow points towards centre. (c) Microtrichia are largely solid cuticular structures (arrowhead), each connected to a fan-fibre (*). (d) Fan-fibres extend to the radial beams, which alternate between thin (blue) and wide beams (red). (e) *In vivo* light microscopy shows the radial beam (*rb*) and the palisade layer (*pl*). (f) Micro-CT confirms that radial beams originate from the dorsoventral palisade layer. Centre-to-centre spacing of the beams is around 4 µm or 1.3°. *vn*: V-notch. Scale bars: (a) 10 µm; (b) 5 µm; (c) 2 µm; (d) 6 µm; (e) 20 µm; (f) 40 µm.

### Attachment performance of blepharicerid larvae on different substrates

#### Effect of surface roughness on the attachment performance of blepharicerid larvae

*H. lugubris* attachment forces were measured using a centrifuge force tester on smooth, micro-rough, and coarse-rough surfaces (surface profiles shown in Table 1). Peak shear and adhesion forces per body weight were measured on horizontal and vertical substrates, respectively (Figure 3a & b). The test substrate had a significant effect on the peak shear force per body weight, with the larvae attaching best on smooth, followed by micro-rough, then coarse-rough substrates (Kruskal-Wallis rank sum test, χ^2^_df=2_= 26.3, *p* < 0.001; *p* < 0.05 for all pair-wise comparisons using Dunn’s *post hoc* tests with Bonferroni-Holm corrections).The same effect was observed for peak adhesion force per body weight (Kruskal-Wallis rank sum test, χ^2^_df=2_= 24.2, *p* < 0.001; *p* < 0.05 for all pair-wise comparisons, see above).

**Table 1.**
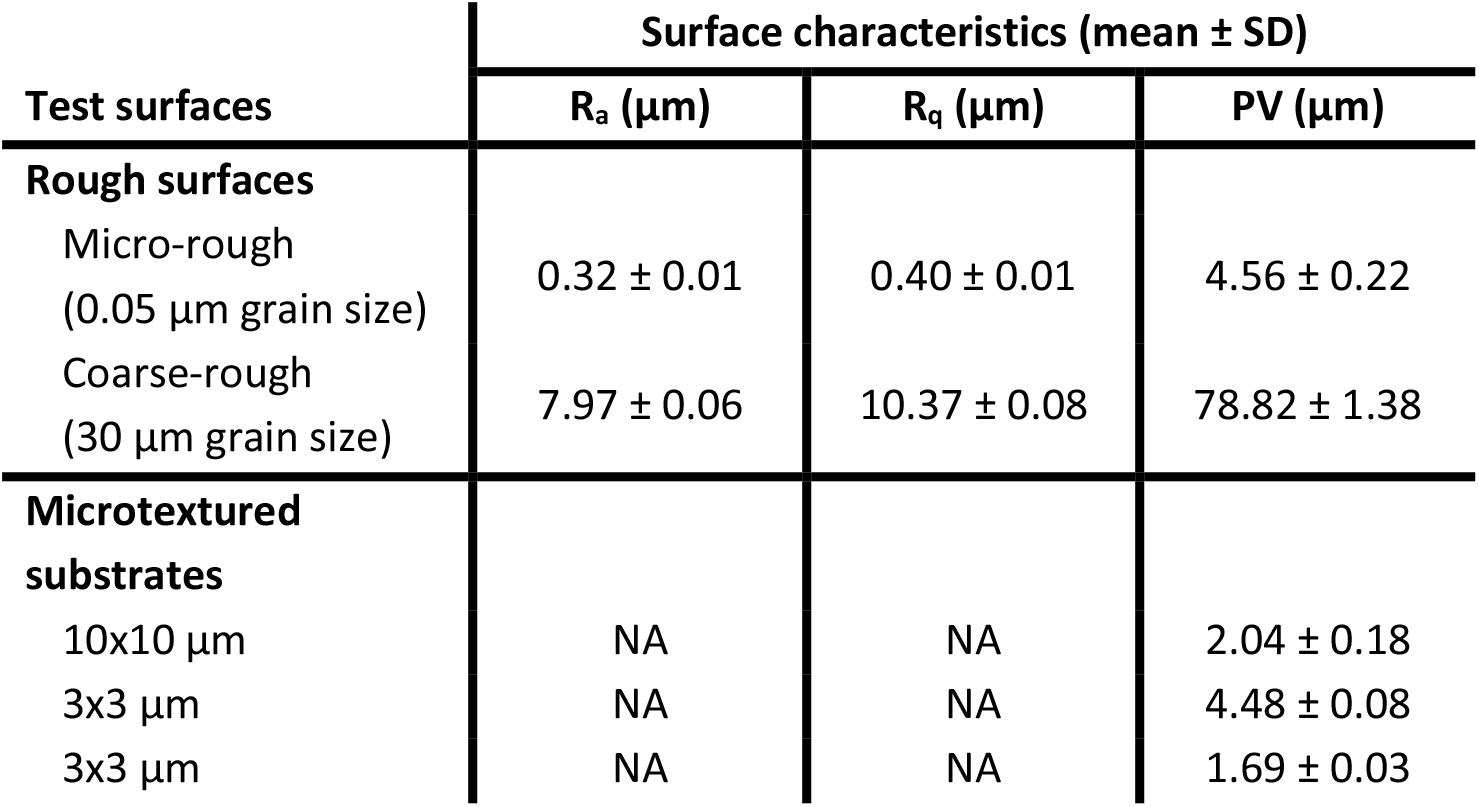
Surface profilometry of test substrates used to assess attachment performance. R_a_: average roughness (mean height deviation), R_q_: root-mean-squared roughness, PV: maximum peak-to-valley height. NA: Not applicable.

**Figure 3.**
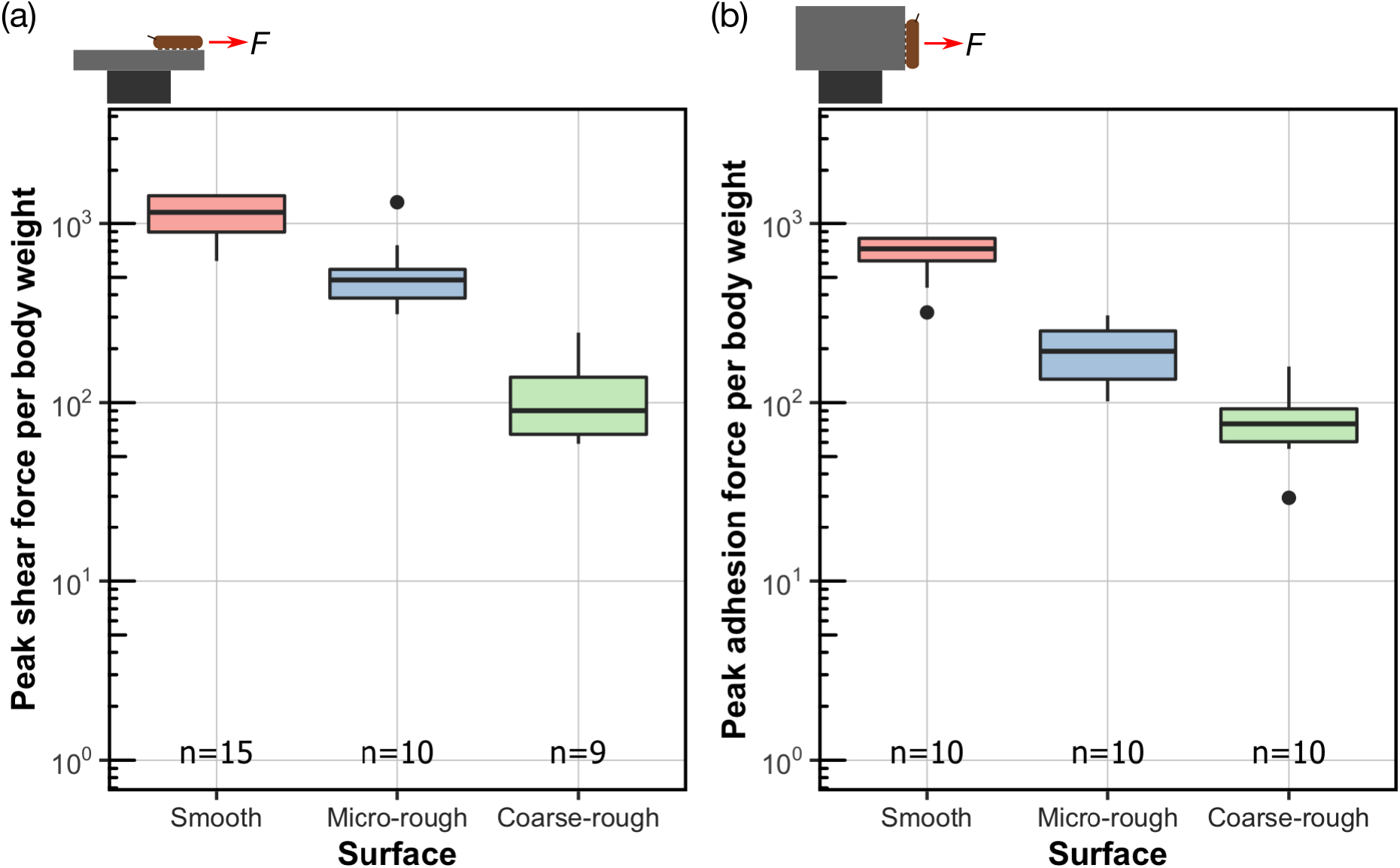
Attachment performance of *H. lugubris* larvae on surfaces of varying roughness. *H. lugubris* larvae performance in (a) peak shear force per body weight and (b) peak adhesion force per body weight.

#### Attachment performance of three blepharicerid species on smooth surfaces

The peak shear force per body weight on smooth surfaces measured for larvae from three blepharicerid species -*L. cordata, L. cinerascens*, and *H. lugubris* - was 585 ± 330, 324 ± 153, and 1,120 ± 282, respectively (mean ± SD; Figure 4a). The highest overall shear force per body weight (1,430) was obtained from a *H. lugubris* larva, while *L. cordata* produced the highest overall shear force of 54 mN.

**Figure 4.**
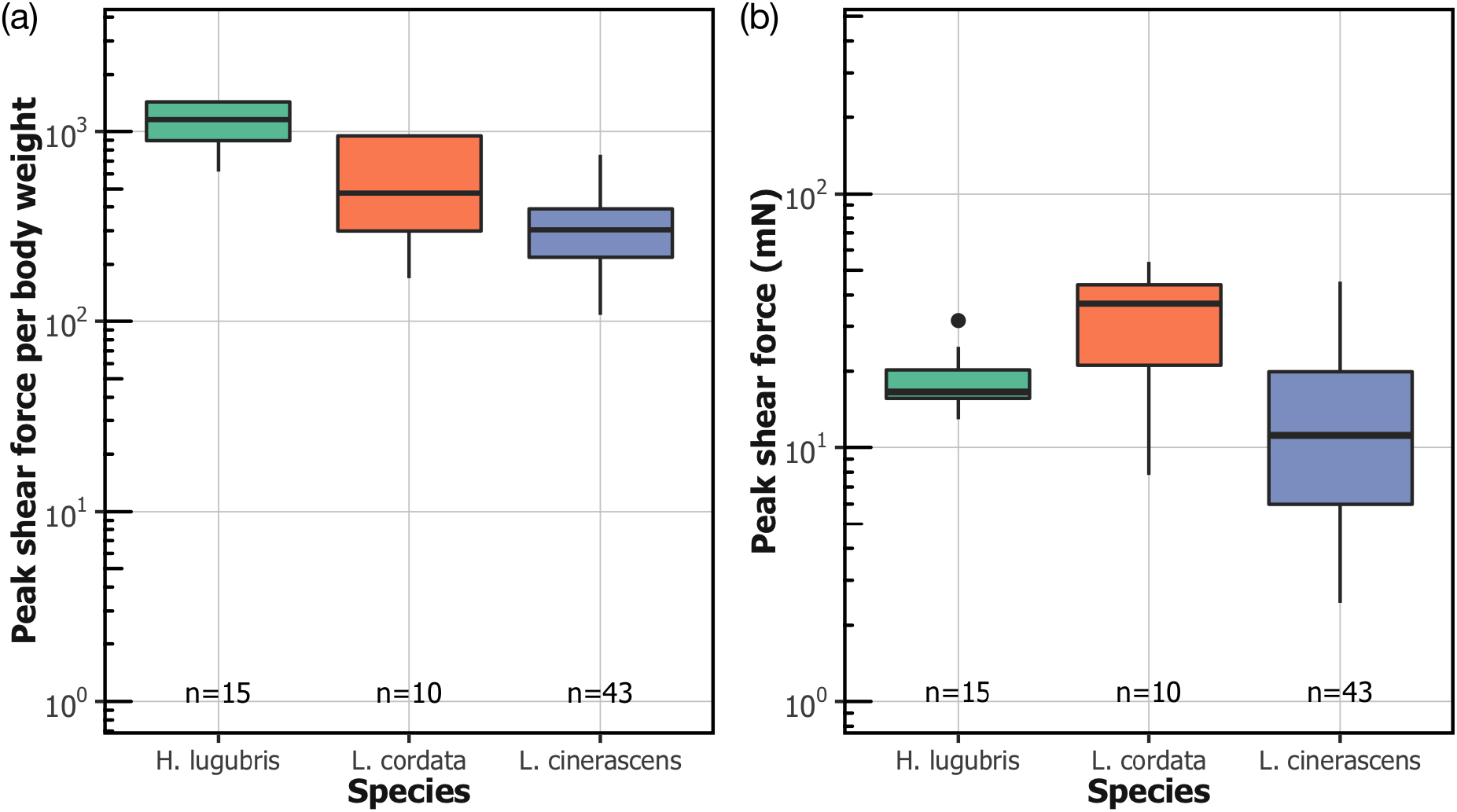
Attachment performance of three species of blepharicerid larvae (*H. lugubris, L. cordata, L. cinerascens*) on smooth horizontal surface. (a) Peak shear force per body weight. (b) Peak shear force. Centre lines, boxes, whiskers, and filled dots represent the median, the inter-quartile range (IQR), 1.5-times IQR, and outliers, respectively.

#### Estimates of peak shear stress on smooth surfaces

Suction disc areas measured for *L. cordata* and *H. lugubris* were used to estimate the peak shear stress for blepharicerid suction attachments. Shear stresses were 41.2 ± 21.4 kPa and 39.3 ± 10.6 kPa (mean ± SD) for *L. cordata* and *H. lugubris*, respectively (Table 2). These values, however, are conservative estimates because: (1) the contact area measurements included the outer fringe which lies outside the suction disc seal, and (2) we assumed that all six suction organs were in contact immediately before detachment. We thus derived more realistic estimates of the shear stresses by first correcting for the fact that the outer fringe layer amounted to 33% to the total imaged contact area (n=18 suction discs from 6 individuals), and second, as larvae attach with fewer suction organs prior to detachment [12]), we assumed that three organs were in contact. Factoring in these assumptions, the shear stresses were 111 ± 57.5 kPa and 117 ± 31.4 kPa (mean ± SD) for *L. cordata* and *H. lugubris*, respectively (Table 2).

**Table 2.**
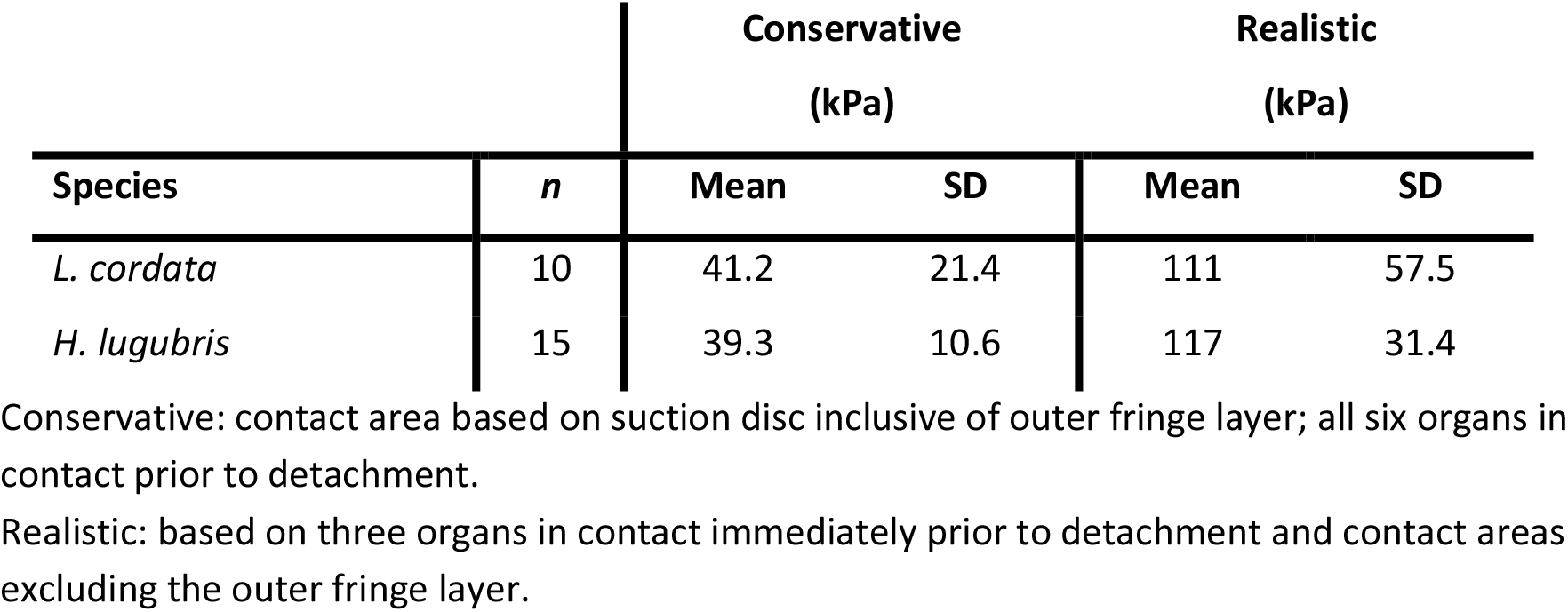
Shear stress estimates for suction-based attachments of *H. lugubris* and *L. cordata*.

#### Shear attachment performance of suction organs on rough substrates compared to smooth adhesive pads

While blepharicerid attachment performance decreased with increasing surface roughness, we observed a different pattern with a model terrestrial climbing insect, *Carausius morosus* stick insects (Figure 5). Stick insects rely on a combination of smooth adhesive pads and claws for attachment, where the former facilitates strong adhesion on smooth surfaces and the latter to coarse-rough substrates. Accordingly, we found that stick insects attached equally well to smooth and coarse-rough surfaces (one-way ANOVA, F_2,27_= 77.0, *p*=0.97 using Tukey’s *post hoc* test; all tests with stick insects were conducted on dry substrates.). On micro-rough surfaces, however, where neither the smooth pad nor the claws proved effective, their shear force per body weight decreased 16-fold (based on the mean of back-transformed values) compared to the smooth surface (same ANOVA as above, *p*<0.001 using Tukey’s *post hoc* test). The attachment performance of *H. lugubris* larvae was also affected by micro-roughness, but to a much lesser degree than in stick insects, with a two-fold decrease (based on the mean of back-transformed values) in shear force per body weight. It is important to mention that blepharicerid larvae do not possess any claw-like appendages that can be used to increase grip.

**Figure 5.**
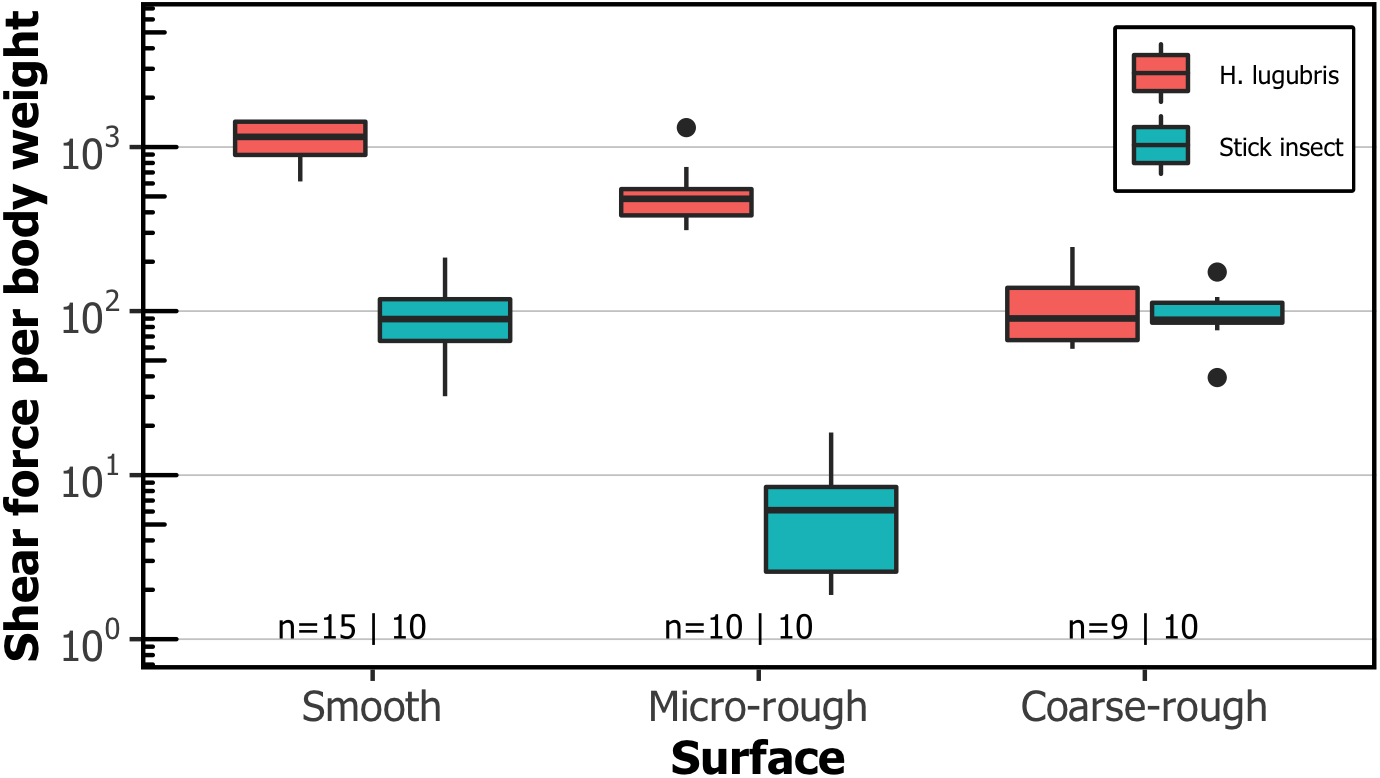
Comparison of shear attachment performance of *H. lugubris* larvae versus stick insects (*Carausius morosus*) on smooth and rough surfaces. *H. lugubris* larvae attach using suction organs, whereas stick insects rely on smooth adhesive pads and claws. Sample sizes shown with *H. lugubris* on the left and stick insects on the right. Centre lines, boxes, whiskers, and filled dots represent the median, the inter-quartile range (IQR), 1.5-times IQR, and outliers, respectively.

### *In vivo* visualisation of suction organs attaching to smooth and transparent microstructured substrates

The contact behaviour of *H. lugubris* suction discs was visualised *in vivo* using interference reflection microscopy (IRM). The attachment-detachment behaviour on smooth glass resembled closely that of the related *Liponeura* species [10, 12]: the suction disc came into close contact with the surface at the outer fringe layer, disc rim, microtrichia zone, and around the central opening (Figure 6a; Video 2). When the piston was raised away from the surface, the greater portion of the suction disc came into close contact as a result of the reduced hydrostatic pressure. When the organ was attached, the microtrichia made tip contact with the surface. No side contact was observed even when the suction disc was pulled closer to the surface as a result of the piston being raised. Detachment of the suction organ and forward movement was often preceded by an active opening of the V-notch (Video 2).

**Figure 6.**
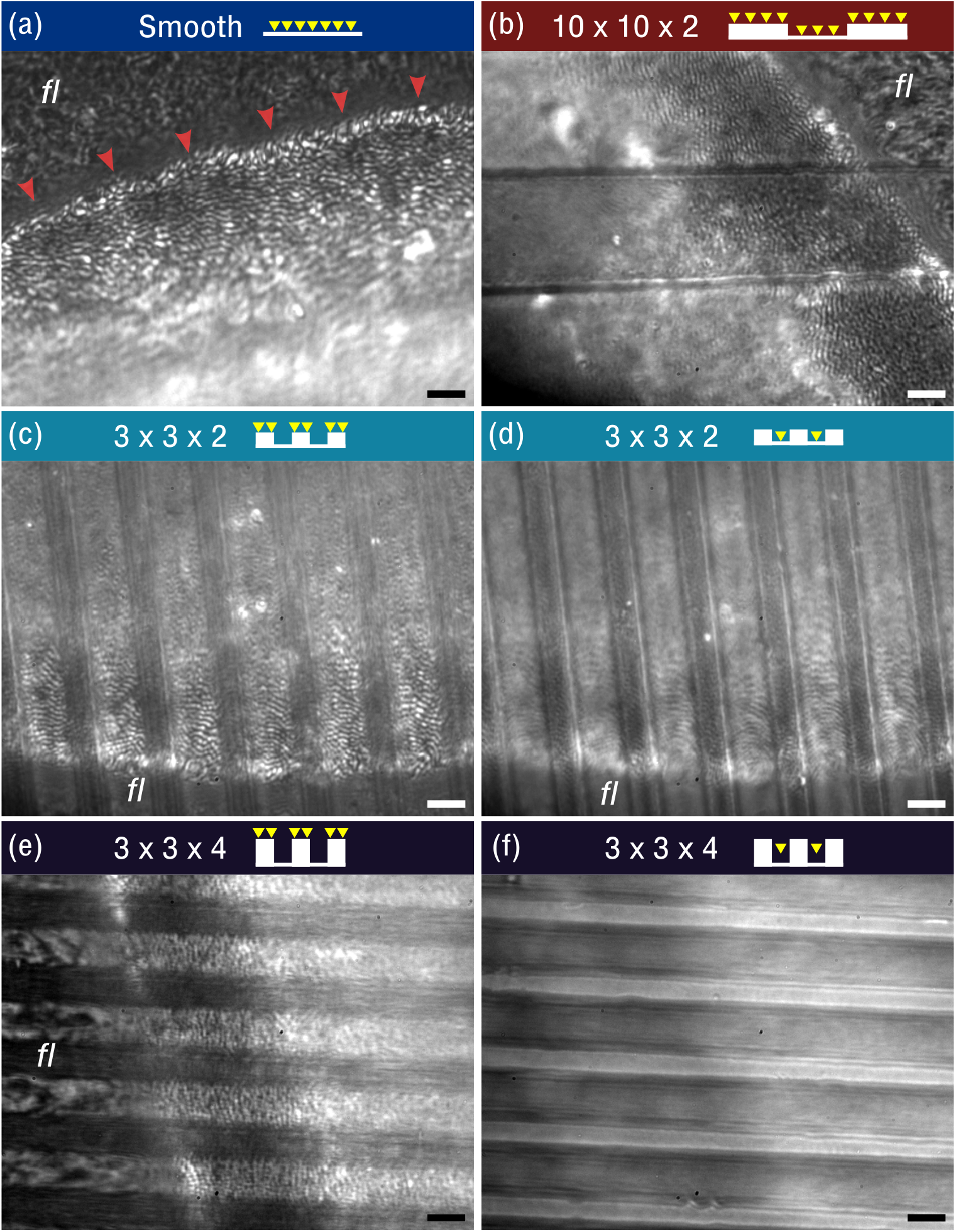
*In vivo* visualisation of *H. lugubris* suction disc contact on different substrates. (a) On smooth glass, microtrichia make tip contact (black dots). Outer fringe layer (*fl*) is outside the seal (red arrowheads). Schematic of the contacts shown with yellow arrowheads representing microtrichia. (b) On 10×10×2 µm microstructured surface (ridges and grooves 10 µm in width, grooves 2 µm deep), contact from microtrichia and *fl* are similar to the contact on the smooth surface. (c) On 3×3×2 µm, the microtrichia made tip contact on the ridges, as well as in the grooves, as seen in (d). However, fewer microtrichia make contact within the narrow grooves compared to the 10×10×2 µm surface. Note: (c) and (d) differ only in the focus settings. (e) On 3×3×4 µm, microtrichia make close contact on the ridges, but inside the deep grooves there is no contact. Scale bars: 3 µm for all images.

Although the outer fringe layer and the microtrichia made close contact on smooth substrates, the outcomes were different on transparent microstructured substrates made of epoxy [26]. On the 10×10×2 µm and 3×3×2 µm substrates (ridge width × groove width × ridge height), the microtrichia made contact on both the ridges and the grooves, which are visible as black dots in the IRM recordings (Figure 6b-d). Similar to our observations on smooth glass surfaces, only the tips of the microtrichia made contact on top of the ridges and inside the grooves. The outer fringe layer also made contact, although not uniformly. In contrast, on the 3×3×4 µm substrate, the microtrichia and the outer fringe layer made contact only on the ridges but not inside the grooves (Figure 6e & f). Moreover, we observed microbial organisms freely floating and moving within the 4 µm deep grooves (confirming the lack of close contact) but not on the ridges where the microtrichia and fringe layer were close to the surface. In contrast, no particulate or microbial movement was observed during the trials with the other microstructured surfaces (3×3×2 µm and 10×10×2 µm).

## Discussion

### Blepharicerid larvae attach with extreme strength on diverse surfaces

Blepharicerid larvae possess some of the most powerful (in terms of body weight) and complex suction organs among animals. The three species of blepharicerid larvae studied here (*H. lugubris, L. cordata*, and *L. cinerascens*) produced extreme shear forces on smooth surfaces with averages ranging from 320 to 1,120 times their own body weight. In terms of weight-specific attachment performance, the larvae perform better than all terrestrial insects measured using comparable methods (i.e., whole-animal detachment experiments) [27, 28]. For example, the weight-specific shear attachment of blepharicerid larvae was 3 to 11 times greater than that of stick insects measured in this study. To achieve this extreme shear attachment on smooth surfaces, blepharicerid suction organs must come in close contact by generating an effective seal. Based on our *in vivo* visualisations of *H. lugubris* attaching to smooth glass underwater, the microtrichia make close contact with the surface, helping to both seal the organ and generate friction. This corroborates our previous findings on the suction disc contact behaviour with *L. cinerascens* and *L. cordata* [10]. Likewise, the soft adhesive pads of stick insects make close contact on smooth surfaces, and while the weight-specific attachment forces are not as high as in blepharicerid larvae, they can withstand forces close to 100 times their body weight.

In contrast, the attachment of stick insects on micro-rough surfaces is significantly different to that of blepharicerid larvae: for stick insects, there was a 16-fold decrease in performance compared to smooth substrates, while for blepharicerid larvae the decrease was only two-fold. This difference in the impact of micro-roughness can be attributed to the two fundamentally different mechanisms of attachment: on micro-rough surfaces, neither the soft adhesive pads nor the tarsal claws of stick insects function properly [29]. This is in part due to the reduced effective contact area (as the adhesive pads cannot mould sufficiently to the asperities), and also due to the reduced friction from tarsal claws (because the claws cannot interlock with the small asperities). On the other hand, blepharicerid suction organs are still able to seal on micro-rough surfaces and microtrichia can interact with the asperities, which likely explains which likely explains why they were not impacted as severely as stick insects. In addition, blepharicerid suction organs may adhere better to micro-rough surfaces because when they are in partial contact, the gaps between the detached regions and the substrate are filled with water, whereas detached regions of stick insect pads are filled with air. As water is effectively incompressible and approximately 50 times more viscous than air, the water-filled contact zone can provide a much stronger resistance against detachment even under conditions of partial contact as on micro-rough substrates.

While blepharicerid larvae attached more strongly than stick insects on micro-rough surfaces, the opposite was found on coarse-rough surfaces: for stick insects, there was no difference in performance between coarse-rough and smooth surfaces, whereas blepharicerid larvae attachment decreased 11-fold. It is likely that both blepharicerid suction organs and stick insect adhesive pads are unable to cope with coarse surface roughness. Stick insects, however, have large pretarsal claws that can interlock with large asperities for strong attachment. Previous studies on dock beetles (*Gastrophysa viridula*) and stick insects *(C. morosus*) reported that both beetle and stick insect attachments on coarse-rough surfaces decrease significantly when the claws are removed [29, 30]. This means that, although stick insects and dock beetles use two distinct adhesive systems (smooth versus hairy pads), the combination of the claws and the adhesive pads produces the same trend: both insects attach strongly to smooth and coarse-rough surfaces but poorly on micro-rough surfaces. In contrast, blepharicerid larvae do not have claw-like appendages and rely on suction organs for attachment; consequently, these aquatic larvae do not follow the same trend as terrestrial climbing insects and perform the worst on coarse-rough substrates. A similar result was reported by Liu *et al*. using *Blepharicera sp*., where the larval attachment performance decreased with increasing surface roughness, although no quantitative information on attachment forces can be extracted from their study [13]. (The study also used a centrifuge, but only reported rotation speed but not the insects’ mass and position; it is also unclear whether the larvae were wetted prior to the tests.)

### Ultrastructural components may help to stabilise the suction disc under high stress

Recent work on cupped microstructures (which resemble microscopic suction cups) has revealed mechanisms of failure in underwater suction attachments: (1) under sustained tensile stress, the rim slides inwards and the rim diameter contracts by ∼30%; (2) immediately prior to detachment, sections of the rim buckle inwards, leading to adhesive failure [31]. Similar failure modes were also reported for macroscopic suction cups [32]. Two structural features of the blepharicerid suction organs could represent adaptations to counter the aforementioned failure mechanisms seen in microscopic suction cups. First, unlike the smooth surface of the fabricated cupped microstructures, blepharicerid suction discs bear numerous microtrichia that can interlock with surface asperities and minimise inward sliding. This has been reported for the remora fish, which have stiff posterior-facing structures called spinules within their suction pads that passively engage with asperities during high-drag conditions [14, 15]; similarly, blepharicerid microtrichia are naturally angled (∼40° relative to the horizontal) and point towards the centre of the suction disc [10]. Hence, inward sliding would passively promote additional interlocking of the microtrichia tips with the surface. Second, the internal radial beams can provide structural support to reduce inward sliding and buckling of the rim. Similar to how the flexible membrane of an umbrella is stiffened by radial spokes, these stiff cuticular radial beams can stabilise the suction disc when the organ is under high tensile stress. Bones within the clingfish suction organ may also prevent inward sliding [32]. While we have yet to visualise blepharicerid suction organs failure under extreme forces, their powerful attachments suggest that they possess mechanisms to counter the common modes of suction cup failure.

### Blepharicerid suction organs are unique among insects and specialised for fast-flow conditions

We have demonstrated that blepharicerid suction organs can attach with extreme strength to both smooth and rough surfaces. Despite the potential strength of each attachment, the larvae are surprisingly mobile in their natural habitat [5, 25]. Both attachment force and mobility are required for blepharicerid larvae to survive in their challenging habitats, which include raging alpine torrents and areas near the base of waterfalls. Currently, blepharicerid suction organs are the only examples of piston-driven suction organs in insects. While the circular setae of male diving beetles (Dytiscidae) are also considered to be suction organs, the two systems have markedly different morphologies: (1) the ventral surfaces of circular setae are comparatively smooth; (2) circular setae lack a muscle-driven central piston; (3) there are no known mechanisms for rapid detachment in dytiscid circular setae [33– 35]. As male dytiscid beetles use their suction organs to attach to the smooth sections of the female’s pronotum and elytra, their attachment works best on smooth surfaces and their performance declines strongly on rough surfaces [34]. Researchers have suggested that male and female dytiscid beetles are engaged in an evolutionary arms race driven by sexual conflict, as female cuticular surfaces are modified to hinder male attachment with suction discs [34, 36]. Interestingly, it appears that male beetles did not evolve friction-enhancing structures on their circular setae to facilitate adhesion to rougher regions of the female cuticle. In contrast, blepharicerid suction discs are covered in a dense array of microtrichia that likely enhance grip on rough surfaces in high-drag conditions. This difference in morphology may be based on function or phylogenetic constraints: for one, male diving beetles may not need to attach to rough elytra if their setae can generate sufficient attachment forces on smooth cuticle alone. Alternatively, beetle setae may be more limited in the structures that can be developed from them, compared to the blepharicerid organ which is highly complex and multicellular.

A more comparable suction-based attachment system can be found in remora fish. Remora use suction pads -highly modified dorsal fin spines - to attach to sharks, whales, and manta rays [14, 15]. Recent studies on the functional morphology of remora suction pads have greatly expanded our understanding on the mechanisms underlying their impressive performance [14, 37, 38]. A remora suction pad comprises a soft fleshy outer rim and rows of lamellae topped with spine-like bony spinules. The pitch of the lamellae is muscle-controlled to facilitate spinule contact with the host skin. When engaged, the tips of these stiff spinules interlock with surface asperities and increase friction, thereby increasing shear resistance. Moreover, the angled posterior-facing lamellae and spinules promote passive engagement when subjected to shear forces from a swimming host [14, 15]. The similarities between the remora suction pad and the blepharicerid suction organ may be based on similar requirements, as both animals have to cope with high shear forces (fast-swimming hosts and torrential rivers for the remora and blepharicerid larvae, respectively). Functionally, the spinules from remora increase shear resistance, allowing the animal to attach strongly to rough sharkskin surfaces. In blepharicerid suction organs, we have shown that microtrichia make tip contact not only on smooth but also on microstructured surfaces. This indicates that the suction organ moulds to surface roughness and microtrichia contact the valleys of these asperities. To avoid buckling and to function effectively, interlocking structures like spinules and claws need to be stiff and strong [38, 39]. Microtrichia are also likely to be stiff structures since (1) they are solid cuticular projections, and dense cuticle can reach high elastic moduli [40]; (2) we only observed tip contact on various surfaces, and even when the disc was pressed again the ridges in microstructured surfaces, we consistently observed tip contact. Hence, the stiff microtrichia interact with rough surfaces and could function in a similar way to remora spinules. Future studies can verify this by measuring the friction coefficient of microtrichia against surfaces with varying roughness.

### Effect of biofilm layer on suction attachments to natural rock surfaces

Since blepharicerid larvae attach to rocks underwater and feed on epilithic algae, their suction organs will in most cases contact biofilm; however, the details of this interaction are unknown [3]. As hypothesised previously, it is possible that the stiff microtrichia penetrate the biofilm layer, [9, 35]. This may allow the microtrichia to directly interlock with asperities on the rock surface or to generate additional friction from embedding numerous microtrichia into the biofilm. Mayfly larvae, which also inhabit fast-flowing watercourses, have friction-enhancing hairs that benefit from interacting with biofilm [41]. Researchers found that a higher proportion of mayfly larvae can withstand fast flow-rates on smooth hard (epoxy) substrates when biofilm is present. Moreover, the setae and spine-like acanthae on the ventral surfaces of mayfly larvae can generate friction forces on clean rough substrates [42]. Additional experiments with blepharicerid larvae are underway to investigate the interaction between stiff microtrichia and soft substrates.

To conclude, we have shown that blepharicerid larvae use their suction organs to generate extreme attachment to diverse surfaces. The suction organ morphology is conserved between *Hapalothrix* and *Liponeura* larvae, comprising of a suction disc that contacts the substrate, dense arrays of microtrichia on the disc surface, muscles to control the piston, and the V-notch detachment system. We also characterised the suction disc ultrastructure, such as internal radial beam structures that could help to stabilise the suction disc when subjected to high stress, and fan-fibres that connect individual microtrichium to the radial beams. In terms of attachment performance, blepharicerid larvae withstand shear forces equivalent to 320 -1,120 times body weight on smooth substrates, depending on the species. *H. lugubris* performed the best overall, reaching weight-specific shear attachments up to 1,430 and estimated shear stresses up to 117 kPa. Although their attachment decreased with increasing surface roughness, blepharicerid suction organs performed better than the smooth adhesive pads of stick insects on micro-rough surfaces. We confirmed that blepharicerid suction organs mould to large surface asperities and that microtrichia come into close contact between the asperities. These microtrichia are stiff spine-like structures that are specialised for maintaining tip contact to the surface for interlocking with asperities. Our study reveals additional mechanistic insights into the function of a highly adapted insect adhesive organ and expands our understanding of biological suction.

## Materials and methods

### Sample collection and maintenance

*Liponeura cinerascens* (Loew, 1845) larvae were collected from fast-flowing alpine rivers near Meiringen, Switzerland (GPS location 46° 44’ 05.6” N, 8° 06’ 55.4” E, in May 2018), and close to Grinzens, Tirol, Austria (47° 12’ 41.4” N, 11° 15’ 28.1” E in September 2018). At the latter site, *Liponeura cordata* (Vimmer, 1916) and *Hapalothrix lugubris* (Loew, 1876) were also collected. For all the species, we collected third and fourth instar larvae that were large enough to be handled for experiments. Wearing fishing waders and diving gloves, we removed rocks from the most turbulent areas of the river and brought them to the riverbank for specimen collection. Although it was previously noted that the larvae can attach so firmly that they are torn upon detachment [43], we found that a gentle nudge using soft-touch tweezers can elicit an evasive response from them, whereupon they could be easily picked up using tweezers and placed in specially prepared 50 mL Falcon tubes. These tubes were first rinsed with stream water, flicked dry to remove most but not all of the water, then a small patch of wet moss was added to the bottom to retain moisture before being sealed with the screw-top cap. All larvae were kept in these tubes in an ice-box during collection and transport. Rocks were returned to their approximate locations after collection.

For long-term maintenance of the larvae, an aquarium tank was set up with water and small rocks from the collection site. A filter unit with two outlets for a small water cascade was used to filter the water and to simulate the natural environment. Multiple air-pumps were also placed close to the aquarium walls to provide ample oxygenation and additional regions with turbulent flow. To promote algal growth, an over-tank LED light was set to a 12h day-night cycle. The aquarium was kept in a 4°C climate room (mean temperature of 3.2 ± 0.9°C; mean ± SD) to replicate alpine stream temperatures.

### Scanning electron microscopy of *Hapalothrix lugubris* suction organs

SEM was used to image fourth instar *H. lugubris* larvae as described previously [10]. In brief, samples fixed in 70% ethanol (v/v) were flash-frozen in liquid ethane cooled with liquid nitrogen and freeze-fractured immediately afterwards with a double-edged razor blade on a cooled aluminium block to obtain longitudinal views. Samples were freeze-dried overnight then carefully mounted on SEM aluminium stubs using carbon tape and silver paint. They were then sputter-coated with 15 nm of iridium and imaged using a field emission SEM (FEI Verios 460).

### X-ray microtomography (micro-CT) of blepharicerid suction organ

One *H. lugubris* fourth instar larva was fixed in 2% paraformaldehyde and 2% glutaraldehyde (v/v) in 0.05 M sodium cacodylate buffer (pH 7.4) for 7 days at 4°C. The larva was then dissected into six pieces -each containing one suction organ - and fixed for an additional day. The samples were then rinsed multiple times in 0.05 M sodium cacodylate buffer followed by deionised water before dehydration through a graded ethanol series: 50%, 75%, 95%, 100% (v/v) and 100% dry ethanol. The dehydrated samples were then critical-point dried using 4 flushes of liquid CO_2_ in a Quorum E3100 critical-point drier.

One critical-point dried suction organ was used for imaging via X-ray microtomography (micro-CT). The sample was mounted on a standard dressmaker’s pin using UV-curable glue then imaged using a lab-based Zeiss Xradia Versa 520 (Carl Zeiss XRM, Pleasanton, CA, USA) X-ray Microscope. The sample was scanned at 0.325 µm/pixel with accelerating X-ray tube voltage of 50 kV and a tube current of 90 µA. A total of 2401 projections collected at 20 sec exposure intervals were used to perform reconstruction using a Zeiss commercial software package (XMReconstructor, Carl Zeiss), utilising a cone-beam reconstruction algorithm based on filtered back-projection. Subsequent 3D volume rendering and segmentations were carried out using Dragonfly v4.0 (Object Research Systems Inc, Montreal, Canada) and Drishti v2.6.5 and v2.7 [44].

### Measuring attachment performance of blepharicerid larvae using a centrifuge

Insect attachment forces were measured using a custom centrifuge set-up described previously [27]. The centrifuge operated on the following principle: a platform with the test substrates and the insect was driven by a brushless motor, and a light barrier sensor was triggered per rotation. This signal was used synchronise image acquisition from a CMOS USB camera (DMK 23UP1300, The Imaging Source Europe GmbH, Bremen, Germany), and image frames and their corresponding times were recorded using StreamPix4 software (NorPix Inc., Montreal, Canada). For safety reasons, the maximum centrifugation speed was limited to approximately 75 rotations per second (rps). Some of the blepharicerid larvae could not be detached even at the maximum speed (n=14 out of 136 measurements); in such cases, we used the maximum acceleration of a successfully detached individual from the given species.

Effect of surface roughness on the peak shear force of blepharicerid larvae was measured on the following substrates: smooth (clean polyester film), micro-rough (polishing film with nominal asperity size of 0.05 µm; Ultra Tec, CA, US), and coarse-rough (30 µm polishing film, Ultra Tec). The same substrate types were used for the adhesion tests, but a polished polymethyl methacrylate (PMMA) surface was used as the smooth substrate. Surface characteristics (average roughness (mean height deviation) R_a_, root-mean-squared roughness R_q_, and maximum peak-to-valley height (PV)) of the micro-rough substrates were obtained using white-light interferometry with a scan area of 0.14 × 0.10 mm (Zygo NewView 200, Zygo Corporation, CT, USA). Micro-rough substrates were sputter-coated with 5 nm of iridium prior to scanning to improve the surface reflectivity. As the coarse-rough substrate could not be adequately imaged via white-light interferometry, we used a Z-stack image focal-depth analysis technique as described elsewhere [45] with a scan area of 0.44 × 0.58 mm. For both surfaces, three regions were selected at random and imaged. Interferometry images were analysed using MetroPro software (Zygo), and a custom MATLAB script was used to reconstruct the surface profile from the Z-stack images (The MathWorks, Inc., MA, United States).

Since *L. cinerascens* and *L. cordata* were difficult to maintain in laboratory conditions, these two species were tested only on smooth horizontal surfaces (n=43 and n=10, respectively). The full range of tests (adhesion and friction on smooth, micro-rough, coarse-rough surfaces) were conducted for *H. lugubris* (n=9 to 15 for shear tests; n=10 for all adhesion tests). Prior to the experiments, individuals were selected from the laboratory aquarium and placed inside 50 mL Falcon tubes as described above. This tube was kept on ice for the duration of the experiment. For each run, a larva was carefully removed from the tube and placed on the test surface. A droplet of water (taken from the aquarium) was used to wash excess debris from the insect, and lab tissue paper was used to wick away excess water without removing all moisture from the larva. These steps were necessary to prime the larvae for the centrifugation trials as they often displayed defensive behaviour from being handled. The larvae adhered and remained still once primed, and between two to four repetitions were performed for each larva. All centrifuge trials were conducted within seven days of collection. After the trials, all the larvae were blot-dried on filter paper and weighed using an analytical balance (1712 MP8, Sartorius GmbH, Göttingen, Germany). Statistical analyses were conducted on log_10_-transformed values using R v3.6.2 run in RStudio v 1.2.5033 [46, 47].

### Calculating peak stress values on smooth horizontal plastic surface

Adhesive stress and shear stress (defined as the peak attachment force divided by the contact area) were calculated using suction disc areas measured for *L. cordata* and *H. lugubris* larvae. Larvae were placed on microscope slides so that the suction organs fully contacted the glass and imaged with a stereomicroscope. Every tested *L. cordata* and *H. lugubris* specimen was imaged. A representative organ was selected from each *L. cordata* and *H. lugubris* larva, and the contact area calculated by fitting a circle inclusive of the outer fringe layer using FIJI [48] (https://imagej.net/Fiji). The peak attachment force was then divided by this contact area to determine the peak stress for each larva.

### Measuring peak shear and normal attachment forces of stick insects

*Carausius morosus* (Sinéty, 1901) stick insects were used as a model for terrestrial insect adhesion, and their attachment on surfaces with varying roughness was measured to compare against blepharicerid larval attachment. Second-instar nymphs with undamaged legs and tarsi were selected for centrifuge adhesion and shear experiments using smooth, micro-rough, and coarse-rough surfaces (n=10 per surface). No adhesion forces could be measured on the micro-rough surface as stick insects failed to hold their body weight during preliminary tests. Before each trial, we checked that the specimen was in contact with the surface using all six legs and that the surface was uncontaminated. Stick insects were oriented with the head facing out, and each individual was tested twice and weighed afterwards. The higher attachment force per individual was used as the peak attachment force.

### *In vivo* observation of suction organs attaching to smooth and micro-patterned substrates

In order to examine blepharicerid larvae locomoting for extended periods of time, a custom flow-chamber was built to imitate the fast-flow conditions of their natural environments (Figure 7). Two aluminium plates (approximately 60 × 100 mm in height×width) each with a rectangular window were used to sandwich an inner chamber made out of polydimethylsiloxane (PDMS; Sylgard 184, Dow Corning, MI, USA). This inner chamber had a lemon-shaped chamber to serve as the observation arena, and an inlet and an outlet for water circulation. Two microscope coverslips (0.16 -0.19 mm thickness, Agar Scientific, Stansted, UK) were used to encase the inner chamber. Two to five larvae were placed on the bottom coverslip, and once the top coverslip was placed over the arena, four clamps were used to squeeze the aluminium plates and coverslips against the PDMS. The soft PDMS moulded closely to the plates and created a water-tight seal. Aquarium water (kept cool in an ice bath) was pumped via a micro-pump (M200S-V, TCS Micropumps Ltd, UK), and the input voltage was controlled by a microprocessor. The flow-rate was controlled by setting an appropriate pump operating voltage. With this flow-chamber, we recorded *H. lugubris* larvae locomotion and the attachment/detachment of suction organs on smooth glass surfaces via interference reflection microscopy (IRM). IRM has been used previously to investigate the contact between animal adhesive organs and the substrate [49, 50]. Videos were recorded using a USB3 CMOS camera (DMK 23UP1300, see above) and IC Capture software (v2.4.642.2631, The Imaging Source GmbH) at 30 frames per second (FPS).

**Figure 7.**
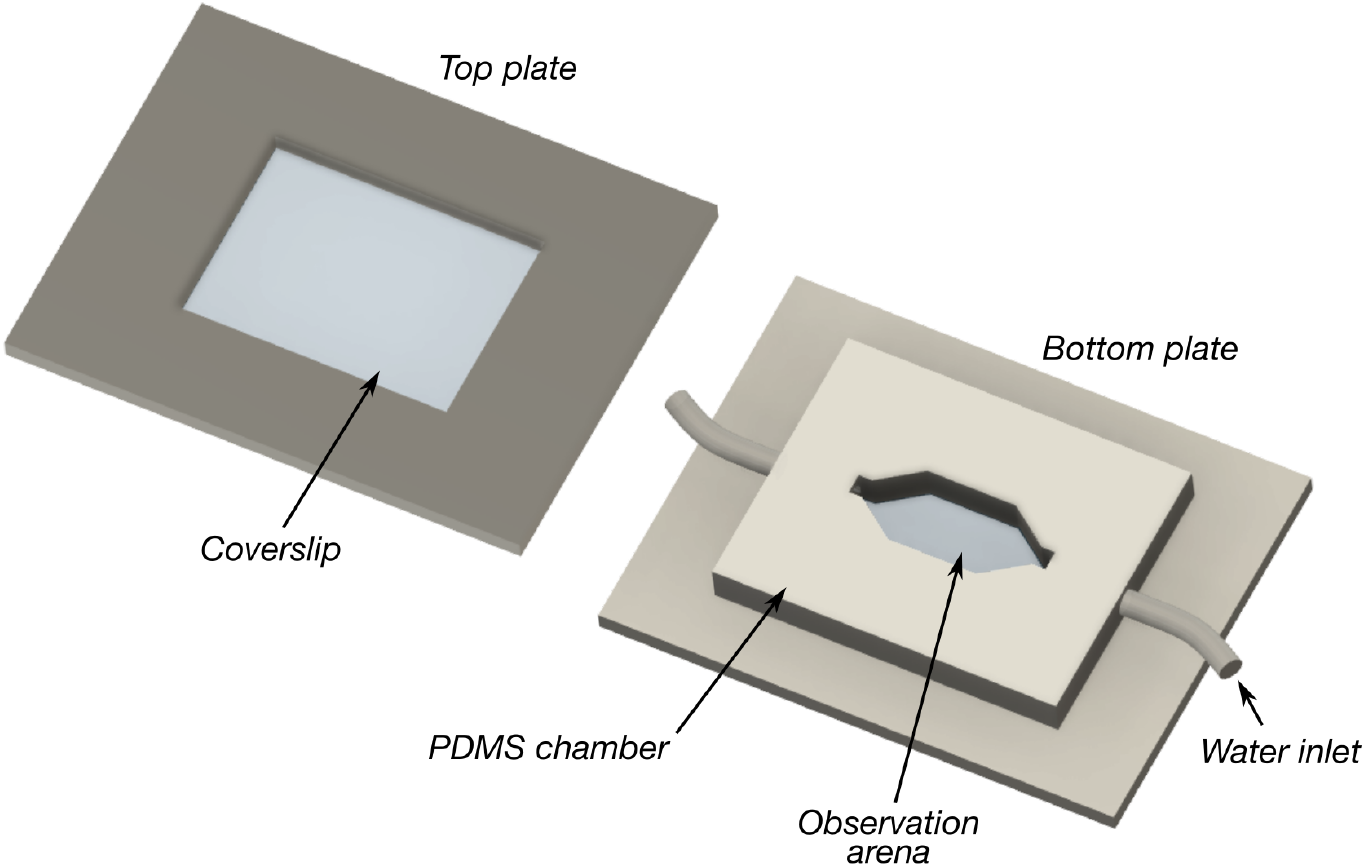
Schematic of the flow-chamber used to observe blepharicerid larvae locomoting in fast-flow conditions. Two to five larvae at a time were added to the observation arena, covered with the top and bottom coverslips and plates, and imaged using interference reflection microscopy. A water pump continuously circulated cooled water at flow-rates ranging between approximately 6 to 15 mL/s. Coverslip thickness: 0.16 -0.19 mm. Plate dimensions: 100 × 60 mm in length×width. Observation arena dimensions: approximately 16 × 8 mm in width×height. PDMS: polydimethylsiloxane

To observe how suction organs respond to surface roughness, we used transparent micro-structured surfaces with well-defined micro-ridges and grooves fabricated by photolithography and nanoimprinting [26]. In brief, a master was first produced using photolithography, and a PDMS mould of this master was then used to cast the final surface out of epoxy. Three micro-ridge geometries were used in our experiments: (1) 3×3×2 µm (ridge width × groove width × ridge height); (2) 3×3×4 µm; and (3) 10×10×2 µm. As ridge height is only approximately controlled through the spin-coating of photoresist when producing the master, we measured it from the epoxy replicas using white-light interferometry as mentioned above (see Table 1 for the results). Four to five regions from each uncoated substrate were imaged and only regions without artefacts were used in the final calculation.

For simplicity, when referring to the substrates, the depths of the grooves were reported to 1 significant figure (i.e., 3×3×2 µm, 3×3×4 µm, and 10×10×2 µm, for widths of ridges, grooves, and ridge height). Note that as these surfaces could not be used in combination with the flow-chamber, a *H. lugubris* larva was placed on the substrate, wetted with a droplet of aquarium water, gently motivated with soft-touch forceps, and recorded as they moved around on the microstructured surface. One *H. lugubris* specimen was used for both the 3×3×2 µm and 10×10×2 µm substrates, and a different larva was used on the 3×3×4 µm surface.

## Acknowledgements

The authors would like to thank P. Ladurner for assisting in sample collection and providing laboratory space near to the field site and M. Sutcliffe for helping with the surface profilometry measurements. They are also grateful to K.H. Muller and J.N. Skepper at the Cambridge Advanced Imaging Centre for their help in preparing and imaging SEM samples.

Rich Media File captions

Video 1. 3-dimensional rendering of a *Hapalothrix lugubris* suction organ based on computed microtomography data. The video begins with a sagittal view of the organ and its internal structures (see Figure 1c – e). Digital dissections and rendering made using Drishti [44].

Video 2. Suction organ of a *Hapalothrix lugubris* larva in action, filmed using *in vivo* interference reflection microscopy and a custom flow-chamber. Note the V-notch opens immediately prior to detachment.

